# FLN-1/Filamin is required to anchor the actomyosin cytoskeleton and for global organization of sub-cellular organelles in a contractile tissue

**DOI:** 10.1101/2020.07.03.186965

**Authors:** Charlotte A. Kelley, Olivia Triplett, Samyukta Mallick, Kristopher Burkewitz, William B. Mair, Erin J. Cram

## Abstract

Actomyosin networks are organized in space, direction, size, and connectivity to produce coordinated contractions across cells. We use the *C. elegans* spermatheca, a tube composed of contractile myoepithelial cells, to study how actomyosin structures are organized. FLN-1/filamin is required for the formation and stabilization of a regular array of parallel, contractile, actomyosin fibers in this tissue. Loss of *fln-1* results in the detachment of actin fibers from the basal surface, which then accumulate along the cell junctions and are stabilized by spectrin. In addition, actin and myosin are captured at the nucleus by the linker of nucleoskeleton and cytoskeleton complex (LINC) complex, where they form large foci. Nuclear positioning and morphology, distribution of the endoplasmic reticulum and the mitochondrial network are also disrupted. These results demonstrate that filamin is required to prevent large actin bundle formation and detachment, to prevent excess nuclear localization of actin and myosin, and to ensure correct positioning of organelles.

## Introduction

Actin and nonmuscle myosin assemble to form force-generating contractile networks. Nonmuscle cells organize their actomyosin cytoskeletal networks to perceive, respond to, and produce forces (Agarwal & Zaidel-Bar, 2019). For instance, neurons have intricate subcellular longitudinal, branched, and periodic networks within dendritic spines (Bär, Kobler, van Bommel, & Mikhaylova, 2016), and the *Drosophila* trachea contains rings of actin cables along the apical surface of the tissue (Matusek et al., 2006). In order to achieve these varying architectures, cells utilize an array of actin interacting proteins. Actin fibers can be linear or branched depending on nucleation and polymerization factors (Pollard, 2007). Filamentous actin (F-actin) can be crosslinked, bundled, severed, or bind scaffolding proteins (Blanchoin, Boujemaa-Paterski, Sykes, & Plastino, 2014; Pollard, 2016). Nonmuscle myosin activity can also shape and organize the actomyosin network (Blanchoin et al., 2014; Shutova, Yang, Vasiliev, & Svitkina, 2012; Wirshing & Cram, 2017). Together these proteins shape the actin cytoskeleton to meet the needs of each cell. We are interested in understanding how filamin, a large, dimeric, actin crosslinker (Feng & Walsh, 2004; Pollard, 2016) functions *in vivo* and in real time to organize and anchor the contractile actomyosin network.

We use the *Caenorhabditis elegans* spermatheca, a contractile tube consisting of 24 myoepithelial cells (McCarter, Bartlett, Dang, & Schedl, 1999), to study how the cytoskeleton becomes organized in a multicellular tissue (Kelley, Wirshing, Zaidel-Bar, & Cram, 2018; Wirshing & Cram, 2017, 2018). The spermatheca is part of the somatic gonad and is the site of fertilization (Strome, 1986). The hermaphroditic reproductive system is made up of two U-shaped gonad arms that surround the developing oocytes, two spermathecae which store sperm, and a shared common uterus (Strome, 1986). One oocyte at a time is ovulated into the spermatheca (McCarter, Bartlett, Dang, & Schedl, 1997; McCarter et al., 1999), where it is immediately fertilized and the eggshell begins to form (Maruyama et al., 2007). Next, the spermatheca coordinately contracts to push the embryo through the spermathecal-uterine (sp-ut) valve and into the uterus (McCarter et al., 1999).

Coordinated contraction of the spermatheca relies on activation of nonmuscle myosin to drive contraction of the actomyosin network (Kelley et al., 2018; Wirshing & Cram, 2017). This network is highly organized and stereotyped (Wirshing & Cram, 2017), with evenly spaced, basal actin stress fibers oriented in parallel to the long axis of each cell (Strome, 1986; Wirshing & Cram, 2017). Prior to ovulation, the network is disorganized, however during the first ovulation, parallel, basal stress fibers form and are maintained throughout the reproductive lifetime of the worm. Organization of the actin fibers requires myosin activity (Kelley et al., 2018; Wirshing & Cram, 2017). In addition to myosin activity, our lab and others have found that several other proteins, including spectrin (Wirshing & Cram, 2018), filamin (Kovacevic & Cram, 2010), actin-interacting protein 1(K. Ono & Ono, 2014), polarity proteins (Aono, Legouis, Hoose, & Kemphues, 2004), formin-related proteins (Hegsted, Wright, Votra, & Pruyne, 2016), and a gelsolin-related protein (Deng, Xia, Fang, & Zhang, 2007) are required for the development and/or maintenance of the organized actin network in the spermatheca.

Filamins contain an N-terminal actin binding domain, two long rod domains composed of immunoglobin (Ig) repeats and separated hinge domains, and a C-terminal self-association dimerization domain to crosslink actin networks (Gorlin et al., 1990). Filamins interact with many signaling and scaffolding proteins including kinases (Kim et al., 2010; Vadlamudi et al., 2002), ion channels (Petrecca, Miller, & Shrier, 2000; Sharif-Naeini et al., 2009), regulators of transcription (Kircher et al., 2015; Sasaki, Masuda, Ohta, Ikeda, & Watanabe, 2001), receptors (Castoria et al., 2011; Li, Bermak, Wang, & Zhou, 2000; Ozanne et al., 2000), and other cytoskeletal proteins (Kim et al., 2010; Labeit et al., 2006). In addition to organizing actin networks (Feng & Walsh, 2004; Pollard, 2016), filamins provide mechanoprotection and durability to sense and withstand forces (Pinto, Senini, Wang, Kazembe, & McCulloch, 2014; Sharif-Naeini et al., 2009; Shifrin, Arora, Ohta, Calderwood, & McCulloch, 2009), modulate ion signaling (Lopez et al., 2018; Petrecca et al., 2000; Rafizadeh et al., 2014; Sharif-Naeini et al., 2009), affect protein trafficking and stability (Li et al., 2000; Ozanne et al., 2000; Petrecca et al., 2000; Rafizadeh et al., 2014), and regulate cell migration (Castoria et al., 2011; Kircher et al., 2015).

*C. elegans* have two filamin orthologs, FLN-1 and FLN-2 (DeMaso, Kovacevic, Uzun, & Cram, 2011). FLN-1 is expressed in the somatic gonad including the gonadal sheath cells, spermatheca, sp-ut valve, and uterus. FLN-2 is not expressed in the somatic gonad, however, it is expressed in the hypodermis, pharynx, intestine, anal depressor muscle, vulva, and distal tip cell (DeMaso et al., 2011; Kovacevic & Cram, 2010). This allows us to reveal functions of filamin that would otherwise be masked by the redundancies seen in many systems, such as human cell culture, which have multiple isoforms (Feng & Walsh, 2004). We have previously reported that FLN-1 localizes to and is required for the development of an organized actin network in the spermatheca (Kovacevic & Cram, 2010). Without FLN-1, the actin network fails to develop organized fibers and is unable to contract, resulting in “trapping” of oocytes within the spermatheca and a low brood size (Kovacevic & Cram, 2010). Interestingly, the disorganized actin phenotype appeared to worsen over time (Kovacevic & Cram, 2010). To better understand this phenotype, we utilized time-lapse 4D microscopy to image the actin network in *fln-1* animals during ovulation. We find that filamin is required to anchor actin bundles at the basal surface, to maintain the organized evenly spaced stress fibers seen in wild type animals, and to prevent actin and myosin from clumping near the nucleus. We also uncover a role for the linker of nucleoskeleton and cytoskeleton complex (LINC) for organizing the actomyosin cytoskeleton. Without FLN-1, a separate pool of actin, unbound by myosin, forms thick peripheral bundles. Several genes stabilize these bundles, including α-spectrin, SPC-1, and β-spectrin, UNC-70. FLN-1, either through its role in anchoring and organizing the actomyosin cytoskeleton or through a novel mechanism, is required for the organized distribution of nuclei, mitochondria, and the endoplasmic reticulum.

## Results

### Filamin is required for actin organization

Filamin is a well characterized actin crosslinker (DeMaso et al., 2011; Nakamura, Stossel, & Hartwig, 2011; Pollard, 2016; Popowicz, Schleicher, Noegel, & Holak, 2006; Razinia, Mäkelä, Ylänne, & Calderwood, 2012; Zhou, Hartwig, & Akyürek, 2010), and loss of filamin results in a disorganized spermathecal actin network (Kovacevic & Cram, 2010). However, many questions remain about the roles of filamin in anchorage and robustness of the actin cytoskeleton. In wild type animals, the spermathecal cells form parallel, basal actin bundles (Figure 1A), while *fln-1(tm545)* animals develop thick peripheral actin cables with thinner, wispy actin fibers distributed throughout the center of the cell (Figure 1B). To understand the temporal dynamics behind these differences we used 4D confocal microscopy to image first ovulations and observe network development using animals expressing ACT-1∷GFP to label actin. In wild type animals, prior to the first ovulation, actin is disorganized (Figure 1C), but as the tissue contracts, the actin becomes organized into parallel bundles oriented along the long axis of the cell (Image 1C’-C’’’, Supplemental Movie 1). Animals lacking *fln-1* also initially have a poorly organized actin network (Image 1D, Supplemental Movie 2). During ovulation, the network fails to become organized into parallel bundles. The actin fibers are pulled to a single point in the cell, relax, and then over time, thicker bundles form along cell peripheries (Figure 1D’-D’’’, Supplemental Movie 2). These results indicate that filamin is required for the development of organized actin bundles, and without filamin, the network becomes progressively more disorganized.

**Figure 1:**
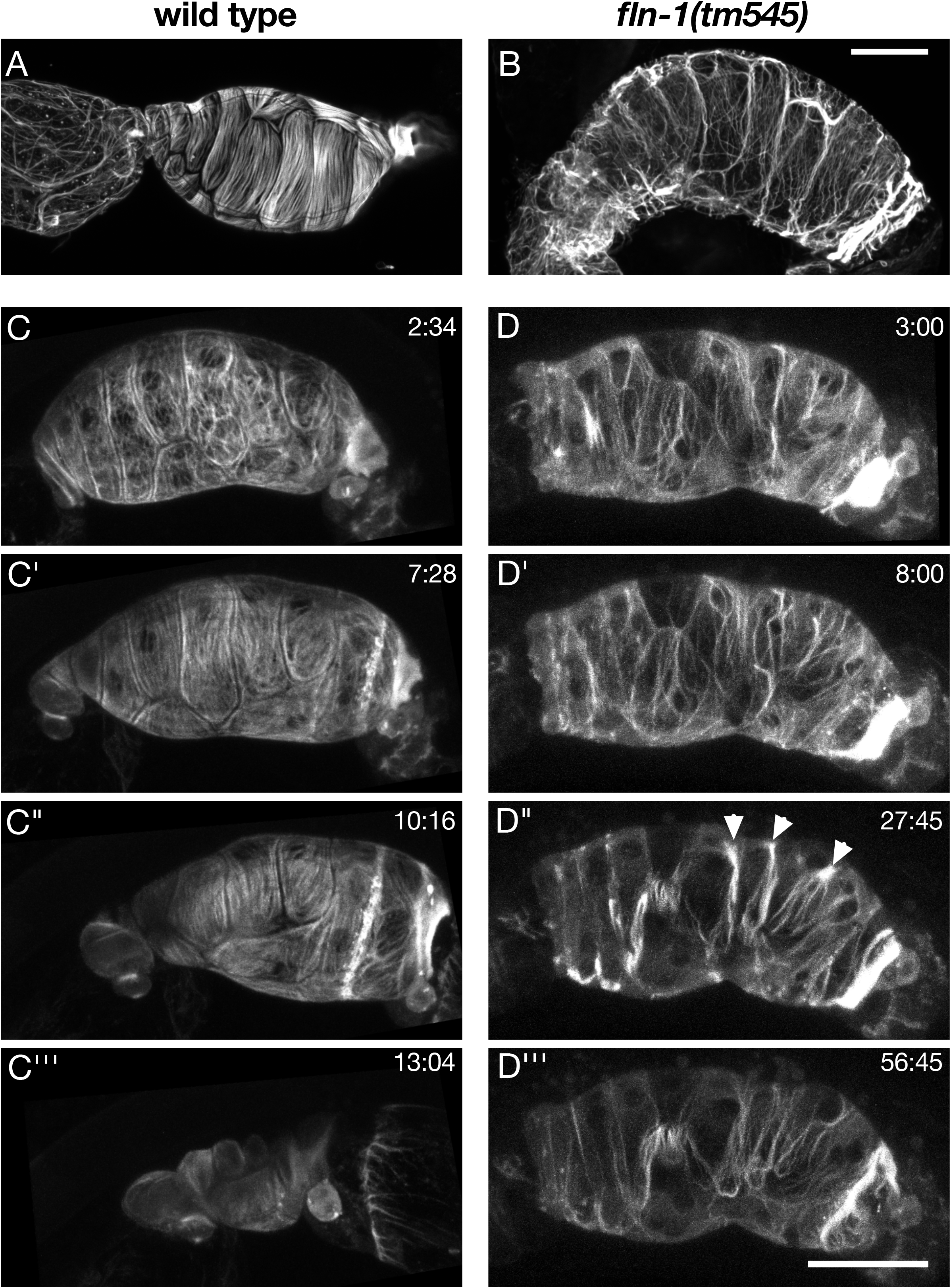
FLN-1 is required for the development of an organized actin in the spermatheca. (A) Phalloidin staining labels the adult wild type spermathecal actin network, which is organized with long parallel actin bundles spanning the length of the cell and oriented along each cell’s long axis. (B) Phalloidin staining of *fln-1* animals reveals disorganized actin networks with thin mesh-like actin fibers within cells and thicker bundles near the cell periphery. (C) Just after oocyte entry of the first ovulation, the actin network in wild type animals, labeled with GFP∷ACT-1, is initially disorganized. (C’) As the sp-ut valve opens, and the spermathecal bag contracts, fibers become aligned and organized as the spermatheca pushes the embryo into the uterus (C’’). (C’’’) The embryo is completely expelled from the spermatheca in wild type animals. (D) In animals lacking *fln-1*, the actin network is also initially disorganized, and remains disorganized through the time when wild type animals have already begun to organize their networks (D’). (D’’) The spermatheca fails to contract and after several minutes, actin briefly forms puncta (arrowheads), before returning to a disorganized, thin meshy network with thicker peripheral bundles (D’’’). Scale bars are 20 μm.

### Myosin forms perinuclear foci in *fln-1* animals

To determine whether myosin activity drives the movement of actin fibers towards the cell center, we imaged first ovulations of animals expressing GFP∷NMY-1 and moeABD∷mCherry to label nonmuscle myosin and filamentous actin, respectively. In these videos, myosin is evenly distributed and colocalizes with actin during contraction (Figure 2A-A’’’, Supplemental Movie 3). Surprisingly, loss of filamin results in the rapid formation of large myosin foci (Figure 2B-B’’’, Supplemental Movie 4). Myosin appears to pull a portion of the actin to the center of the cell, while a population of actin remains near the cell periphery. We quantified colocalization of actin and myosin in wild type and *fln-1* animals starting after the egg is fully inside the spermatheca (Figure 2A’, B’). In wild type animals, actin and myosin briefly become less colocalized after egg entry into the spermatheca, but become more strongly colocalized during contraction and egg exit (Figure 2C). This result is consistent across several experiments (Supplemental Figure 1), and with previous research demonstrating myosin phosphorylation and activation results in an increase in actomyosin colocalization (Kelley, De Henau, Bell, Dansen, & Cram, 2020; Kelley et al., 2018; Wirshing & Cram, 2017). The *fln-1(tm545)* animals exhibit a slow decrease in actomyosin correlation, followed by a precipitate drop as separate foci form (Figure 2B’’), and recovery as the tissue relaxes (Figure 2D). We conclude that filamin is required for the correct distribution and colocalization of the actomyosin network.

**Figure 2:**
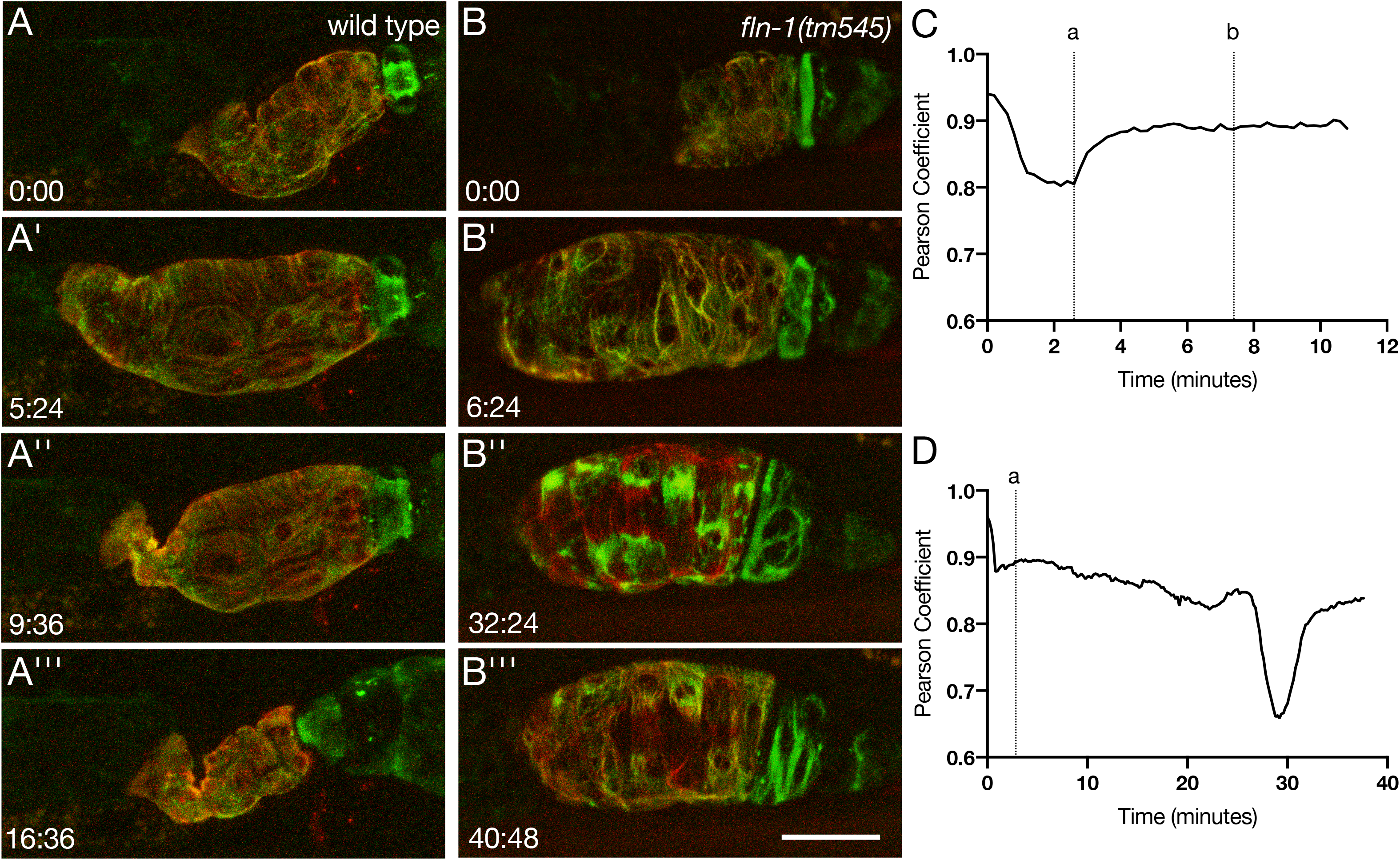
FLN-1 is necessary for colocalization and distribution of actin and myosin. (A-A’’’) Maximum intensity projection of animals expressing moeABD∷mCherry and GFP∷NMY-1 to label F-actin (red) and myosin (green), respectively. Frames show the actomyosin network prior to oocyte entry (A), just after oocyte entry (A’), and the start of embryo exit (A’’), and after ovulation (A’’’). (B-B’) Maximum intensity projection of actin (red) and myosin (green) in *fln-1(tm545)* animals. Frames show the actomyosin network prior to oocyte entry (B), just after oocyte entry (B’), the point at which myosin forms puncta (B’’), and the network after puncta formation (B’’’). Scale bar is 20 μm, both movies were imaged at 12-second intervals. (C) Pearson coefficient of wild type ovulation shown in (A). Correlation is measured from the time point that the egg begins to enter the spermatheca until the end of imaging. The point where the egg is fully entered the spermatheca (A’), and where the egg begins to exit the spermatheca (A’’), is indicated with the dashed lines labeled “a” and “b” respectively. (D) The Pearson correlation coefficient was measured for each frame of the *fln-1(tm545)* ovulation shown in (B), from when the oocyte has begun to enter the spermatheca, until the end of the imaging session. The dashed line labeled “a” indicates when the egg has fully entered the spermatheca (B’).

To more precisely determine the localization of the myosin foci, we excised and fixed gonads of wild type and *fln-1* animals coexpressing GFP∷NMY-1 and moeABD∷mCherry to label myosin and actin, respectively, and stained the nuclei using 4’,6-diamidino-2-phenylindolea (DAPI). We found that in *fln-1* animals, myosin foci are always associated with a nucleus (Figure 3A-B). Line scans through the *fln-1* foci indicate that the foci include both actin and myosin (Figure 3C). In contrast, the peripheral actin bundles lack myosin (Figure 3C’). Since GFP∷NMY-1 labels only the heavy chain of myosin (Piekny, Johnson, Cham, & Mains, 2003; Wirshing & Cram, 2017), we asked if these myosin foci were functional nonmuscle myosin hexamers (Conti & Adelstein, 2008). We used labeled myosin regulatory light chain (MRLC), GFP∷MLC-4 (Kelley et al., 2018; K. Ono & Ono, 2016; Shelton, Carter, Ellis, & Bowerman, 1999), and stained F-actin with phalloidin to show that the MRLC is also associated with the nucleus in *fln-1* animals, but not in wild type animals (Figure 3D). We measured the intensity of actin and myosin along an actomyosin fiber associated with the nucleus and found that the intensity of GFP∷MLC-4 lessened as distance from the nucleus increased, while actin intensity remained relatively unchanged (Figure 3D’). These data suggest that loss of filamin results in not only a disorganized actin network but also redistribution of myosin to foci that are in close proximity to the nucleus.

**Figure 3:**
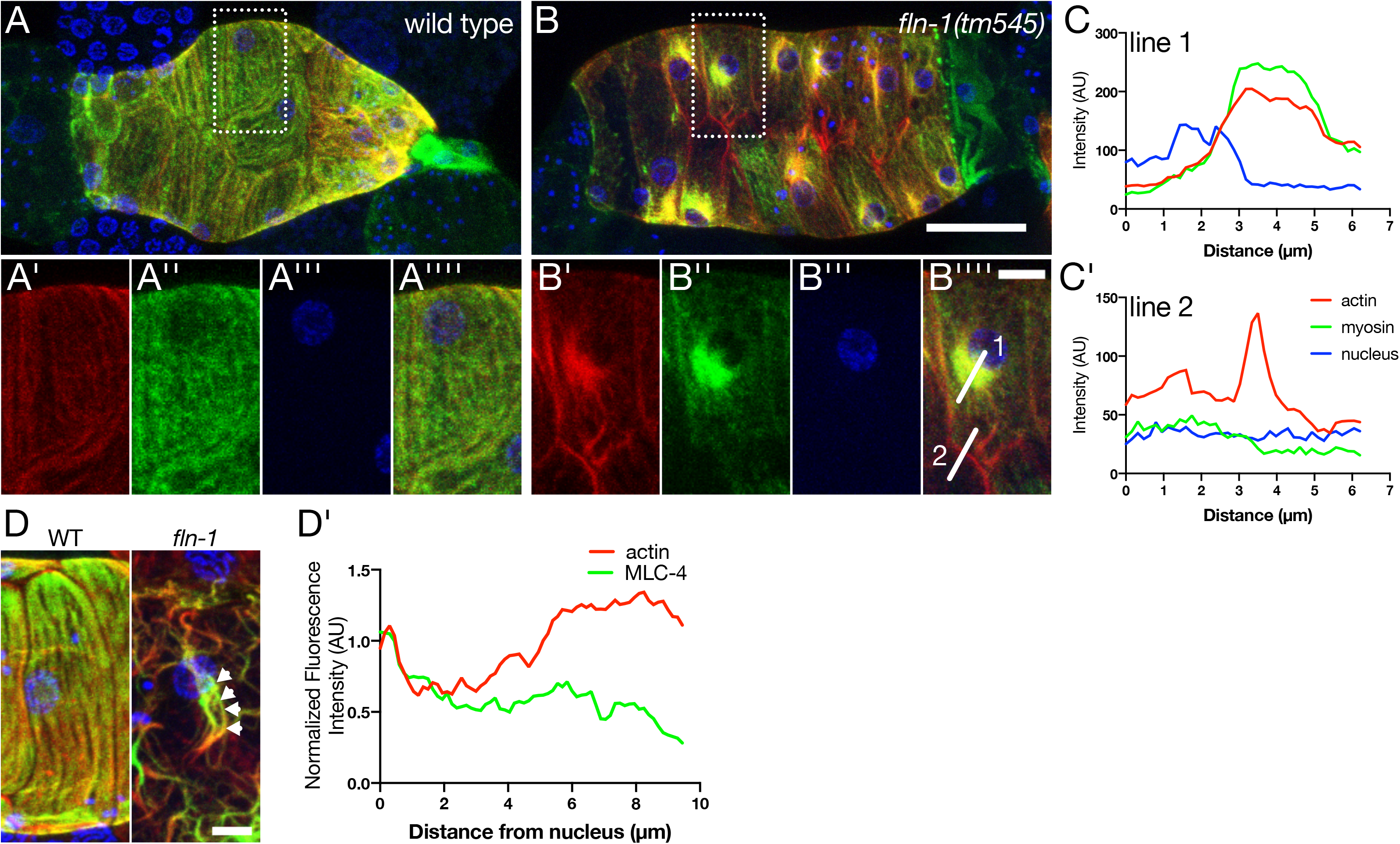
Depletion of *fln-1* results in the formation of perinuclear myosin foci. (A) Maximum intensity projection of excised, fixed, and DAPI (blue) stained spermathecae labeled with moeABD∷mCherry (red) and GFP∷NMY-1 (green). The inset labeled in B shows that actin (A’) and myosin (A’’) are evenly distributed throughout the cell and not clumped near nuclei (A’’’). The merge of all three channels is shown in A’’’’. (B-B’’’’) In *fln-1(tm545)* animals, myosin foci (B’’) are located adjacent to the nucleus (B’’’). Actin (B’) is both associated with the nucleus and is localized to the cell periphery. In A and B, the scale bar is 20 μm. In A’-A’’’’ and B’-B’’’’ the scale bar is 5 μm. (C) Line scan of the nuclear associated actomyosin annotated as “line 1” in B’’’’ shows that myosin and actin intensity is increased and in close association with the DAPI nuclear signal. (C’) “Line 2”, drawn in B’’’’, shows that peripheral actin does not seem to have myosin associated with it. (D) The myosin regulatory light chain (green) labeled with GFP (MLC-4∷GFP) is distributed along phalloidin-stained actin fibers (red) evenly in wild type animals. In *fln-1* animals MLC-4 is still seen along actin fibers. Scale bar is 5 μm. (D’) Quantification of MLC-4∷GFP along a fiber (arrowheads in D) show that MLC-4 intensity decreases as you measure further from the nucleus, while actin remains relatively unchanged.

### The LINC complex is required for the perinuclear localization of actomyosin foci

We next explored what might localize the actomyosin foci to the nucleus in the absence of FLN-1. Nuclei interact with the cytoskeleton through a well conserved, transluminal protein-protein interaction known as the LINC complex (Starr & Fridolfsson, 2010). We knocked down the SUN domain protein, UNC-84, and the KASH domain protein that interacts with actin, ANC-1, in the *fln-1(tm545)* mutants using RNA interference (RNAi) and imaged spermathecae to determine the actomyosin localization with respect to nuclei. We found that knockdown of neither *anc-1* nor *unc-84* affected the formation of actomyosin foci per se, however, the foci were no longer tightly associated with nuclei (Figure 4A-A”). We measured the distance between the edge of the nucleus to the peak of the GFP∷NMY-1 signal, and found that while *fln-1* myosin foci were adjacent to nuclei (0.72 ± 0.07 μm), knockdown of *anc-1* or *unc-84* resulted in significantly larger distances between them (5.97 ± 0.51 μm and 6.68 ± 0.54 μm, respectively) (Figure 4B,C). Endogenously tagged ANC-1 is expressed in the spermatheca and localizes to the periphery of nuclei (Figure 4D-D’’’). Together these data suggest that the LINC complex functions to tether the myosin foci to the nucleus.

**Figure 4:**
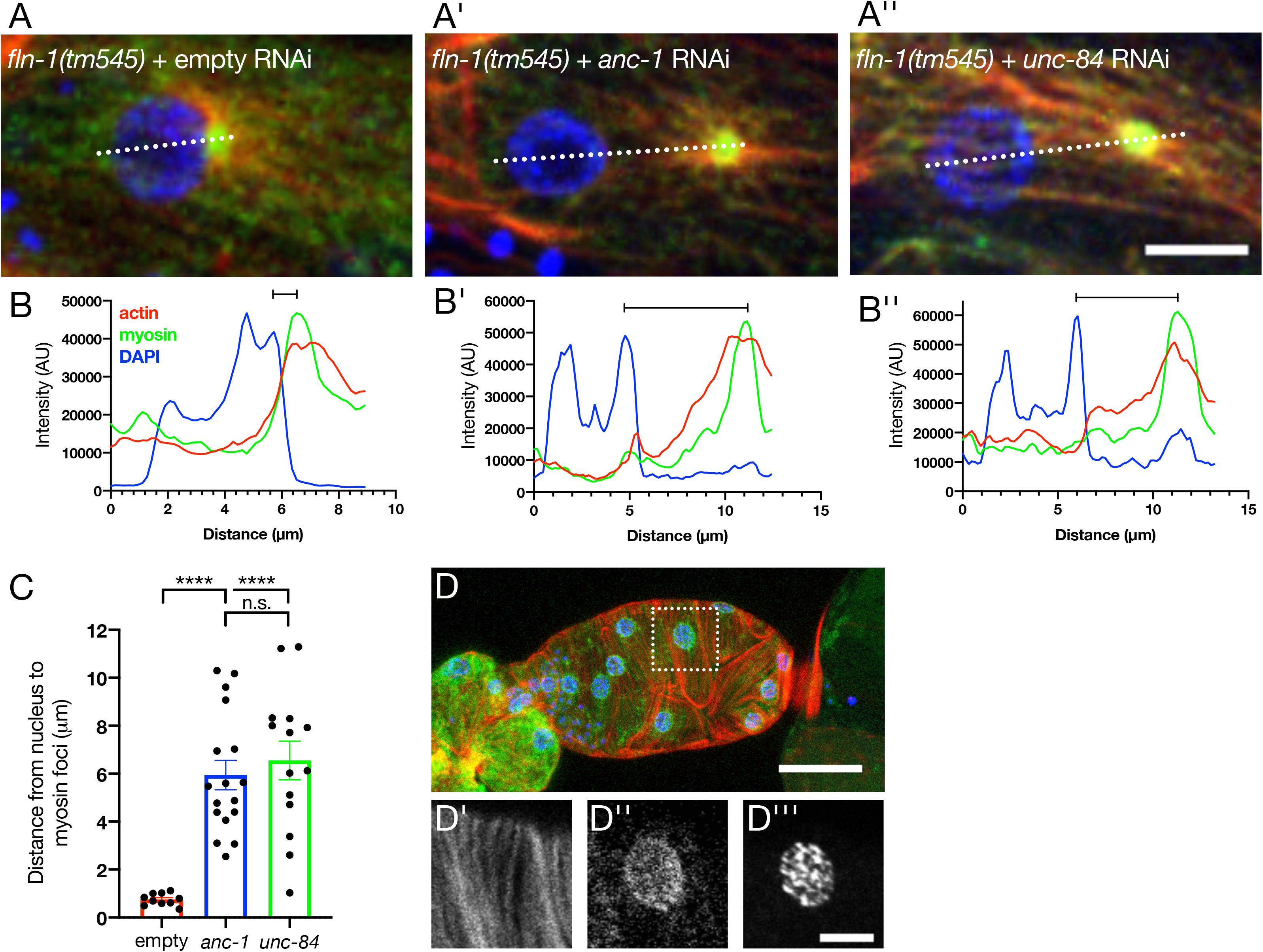
The LINC complex is required for localization of the myosin foci seen in *fln-1* animals. Maximum intensity projections of *fln-1(tm545)* animals treated with either empty (A), *anc-1* (A’) or *unc-84* (A’’) RNAi. Scale bar is 5 μm. Actomyosin puncta are adjacent to the nucleus in empty animals, as visualized by a line scan (B). Quantification (C) shows that distance increases between the nucleus and myosin foci with either *anc-1* (B’) or *unc-84* (B’’) RNAi. Each point represents the average of 2-5 cells per spermatheca. Scale bars are SEM. Ordinary one-way ANOVA with Tukey’s multiple comparisons test: ****p<0.0001. (D) Maximum intensity projection of wild type excised spermatheca with endogenous ANC-1 labeled with a N-terminal GFP tag (green), phalloidin labeled F-actin (red), and DAPI stained nuclei (blue). Inset (dotted box) shows that actin (D’) forms organized, parallel bundles. ANC-1 (D’’) is expressed in the spermatheca and localizes to the nucleus, stained with DAPI (D’’’).

### FLN-1 is required for the basal recruitment and stabilization of the actin network and associated actin-binding proteins

We next investigated differences in the apical-basal polarity of bundle localization in *fln-1* mutants. While wild type animals have contractile actin bundles tethered to the basal surface, in *fln-1* mutants, the majority of the thick actin bundles are displaced apically (Figure 5A-B). Focal adhesions, made up of a complex of proteins including vinculin, connect the basal cytoskeleton to the extracellular matrix (Carisey & Ballestrem, 2011; Golji & Mofrad, 2013; Patel et al., 2006). We examined DEB-1/vinculin distribution by excising gonads of animals expressing DEB-1∷3xGFP and moeABD∷mCherry to label F-actin (Edwards, Demsky, Montague, Weymouth, & Kiehart, 1997; Wirshing & Cram, 2017), and stained the nuclei using DAPI. We found that in wild type animals, DEB-1 decorates the basal actin stress fibers, with some localization to cell junctions (Figure 5C,D). In *fln-1(tm545)* animals, junctional DEB-1 increases compared to the DEB-1 localized to the cell surface (Figure 5 C’,D). In animals lacking *fln-1*, most spermathecae have cells that recruit vinculin to the vicinity of the perinuclear actin (67% N=12; Figure 5E-F).

**Figure 5:**
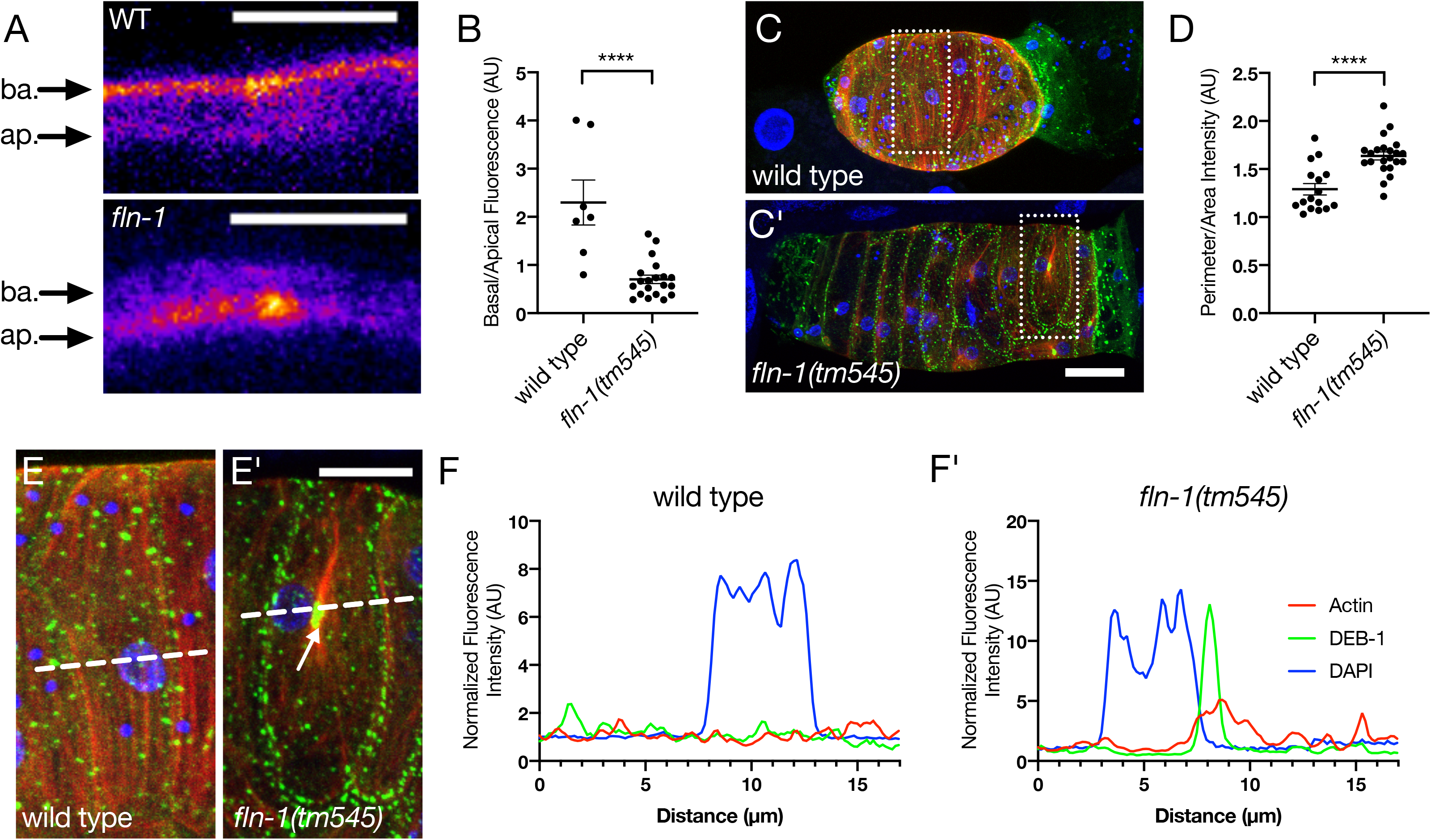
Filamin is involved with actin anchorage to the basal surface. (A) XZ projection through an animal expressing moeABD∷mCherry labeled actin. Fibers are localized more apically in *fln-1* animals compared to the basal fibers seen in wild type. Scale bars 5 μm. (B) Quantification shows that there is a decrease in basal actin fluorescence intensity compared to apical intensity. Each point represents a measurement of a single cell. Maximum intensity projection of excised wild type (C) and *fln-1*(C’) spermathecae where vinculin (green) is labeled with an internal 3xGFP, F-actin (red) is labeled with moeABD∷mCherry, and the nucleus (blue) is stained with DAPI. For A-A’, the scale bar is 20 μm. (D) Peripheral vinculin intensity in *fln-1* animals increases compared to cell surface intensity. Cells (outlined with dotted boxes in C-C’) of wild type (E) and *fln-1*(E’) animals show that there is a redistribution of vinculin upon loss of *fln-1*. Wild type animals have puncta of equal sizes distributed throughout the surface of the cell, with some peripheral signal. *fln-1(tm545)* animals have increased DEB-1 signal along the periphery of cells and near the nucleus (arrow). For B-B’ the scale bar is 10 μm. (F-F’) Line scans (dotted lines annotated in E-E’) show that wild type animals have evenly spaced DEB-1 and actin intensities of relatively equal proportion, and loss of *fln-1* results in a large accumulation of perinuclear DEB-1.

We next asked if this redistribution of DEB-1/vinculin is accompanied by the redistribution of other focal adhesion components such as TLN-1/talin (Cram, Clark, & Schwarzbauer, 2003; Moulder, Huang, Waterston, & Barstead, 1996) and PAT-3/β-integrin (Gettner, Kenyon, & Reichardt, 1995). We knocked down *fln-1* using RNAi in animals expressing endogenously tagged TLN-1 or PAT-3, and stained for F-actin and nuclei using phalloidin and DAPI, respectively. Both proteins are localized broadly across the basal surface of spermathecal cells and colocalize with basal actin fibers, however, in contrast with the gonadal sheath cells, large focal adhesion-like puncta are not observed in the spermatheca (Supplemental Figure 2). With disruption of *fln-1*, we observe localization of TLN-1 and PAT-3 along large actin bundles but no increase in PAT-3 or TLN-1 at perinuclear foci or the cell junctions (Supplemental Figure 2). These data suggest that FLN-1 is required for correct spacing of actin bundles and also functions to keep the fibers at the basal surface and that without filamin, actin and DEB-1 mislocalize to the apical surface, to cell junctions, or to the nuclear periphery.

### Spectrin stabilizes larger, peripheral actin bundles in *fln-1* mutants

Without filamin, the actin network becomes detached from the basal surface, and a portion of the actin pool in *fln-1* mutants is pulled into myosin-rich perinuclear foci. However, a large portion remains along the cell periphery of the cells (Figure 1B). In order to determine the stability of these fibers, we used fluorescence recovery after photobleaching (FRAP) to quantify the turnover of these actin bundles. Since the thicker fibers accumulate near the edges of the cell, we used fiber thickness as a proxy for fiber location in animals where only ACT-1∷GFP was expressed. We bleached both large (>1 μm) and small (≤1 μm) diameter fibers, and found that large fibers had a smaller mobile fraction than the thinner fibers, suggesting increased stability of the large fibers (Figure 6A-C). There was no difference in half-recovery time between fiber categories (Figure 6D). Together this suggests that there is a larger immobile fraction in the thicker fibers, perhaps due to a secondary actin bundling protein that normally competes with filamin, but now is free to bind and stabilize actin fibers.

**Figure 6:**
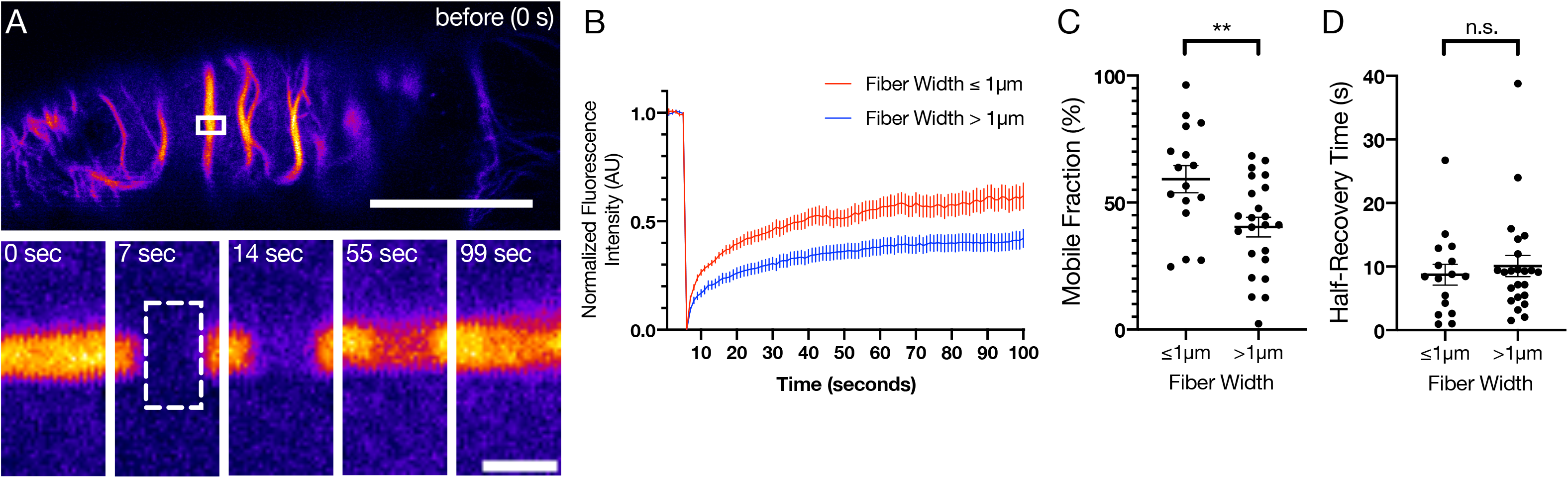
Larger peripheral actin bundles are more stable than smaller bundles. The actin bundles in *fln-1* animals labeled with ACT-1∷GFP were photobleached and fluorescence recovery was monitored over 100 seconds at 1 frame per second. Fiber size was quantified and fibers were grouped into small (≤1 μm) and large (>1 μm) fibers. (A) Representative image of ACT-1 intensity before being bleached (0-seconds), after bleaching (7-seconds), and during recovery (14-, 55-, and 99-seconds). Inset images have been rotated 90° from A. Scale bar is 20 μm and 2 μm for the large and inset images, respectively. (B) Normalized fluorescence intensity of the bleach and recovery curves for large and small fibers in *fln-1* animals. Error bars are SEM. (C,D) The mobile fraction was significantly smaller in large fibers (>1 μm) than in small fibers (≤1 μm), but half recovery time remained unchanged. Each point represents a single bleach and recovery curve. Unpaired t test: ** p ≤ 0.01.

We hypothesized that in the absence of FLN-1, the peripheral actin fibers that detach from the basal surface may be bundled by another actin binding protein, contributing to their increased stability (Figure 6A-D). To identify the protein(s) responsible for the increased bundling, we performed FRAP on *fln-1* mutants treated with RNAi for candidate genes. Our candidates were chosen based on their expression patterns, actin binding or bundling capacity, and their apical localization (Supplementary Table 1). We found that FHOD-1/formin, GSNL-1/gelsolin, PLST-1/plastin, SPC-1/α-spectrin, and VILN-1/villin influence the turnover of actin measured by the mobile fraction (Supplemental Figure 3, Table 1).

**Table 1.**

| Gene | Protein | Mobile Fraction<br>Fibers > 1 μm<br>(% ± SEM) | Mobile Fraction<br>Fibers ≤ 1 μm<br>(% ± SEM) |
| --- | --- | --- | --- |
| empty | | 33.4 ± 3.6 | 55.4 ± 5.4 |
| *alp-1* | ALP/Enigma | 41.9 ± 8.0 | 62.2 ± 10.9 |
| *atn-1* | alpha actinin | 35.7 ± 5.9 | 54.6 ± 5.8 |
| *fhod-1* | formin | 50.2 ± 7.8 | 74.1 ± 2.8 |
| *fli-1* | flightless I | 34.6 ± 5.5 | 51.8 ± 9.0 |
| *gsnl-1* | gelsolin | 49.5 ± 4.8 | 42.6 ± 4.7 |
| *inft-2* | formin | 28.3 ± 6.8 | 43.3 ± 5.4 |
| *plst-1* | plastin | 49.9 ± 6.6 | 68.3 ± 7.9 |
| *spc-1* | alpha spectrin | 53.2 ± 3.2 | 68.3 ± 4.5 |
| *unc-44* | ankyrin | 46.3 ± 9.0 | 60.2 ± 6.7 |
| *unc-115* | | 31.1 ± 4.6 | 44.7 ± 7.0 |
| *viln-1* | supervillin | 38.8 ± 7.0 | 39.0 ± 5.3 |
| *zyx-1* | zyxin | 39.9 ± 5.3 | 60.2 ± 3.7 |

We next further investigated the role of spectrin in stabilizing the large actin bundles seen in *fln-1* animals. Spectrin functions as a tetramer of two alpha spectrin and two beta spectrin subunits oriented in an anti-parallel manner, and localizes to and stabilizes cortical actin networks (Machnicka et al., 2014). We have previously reported that UNC-70 is the β-spectrin subunit expressed in the spermatheca that helps localize SPC-1 to organize the spermathecal actin network (Wirshing & Cram, 2018). Knockdown of *unc-70* recapitulates the result we see in *spc-1* knockdown, with similar mobile fractions of large fibers (Figure 7A-B). No differences in mobile fraction of small (≤1 μm diameter) fibers between control RNAi and knockdown of *spc-1* or *unc-70* were observed (Figure 7B).

**Figure 7:**
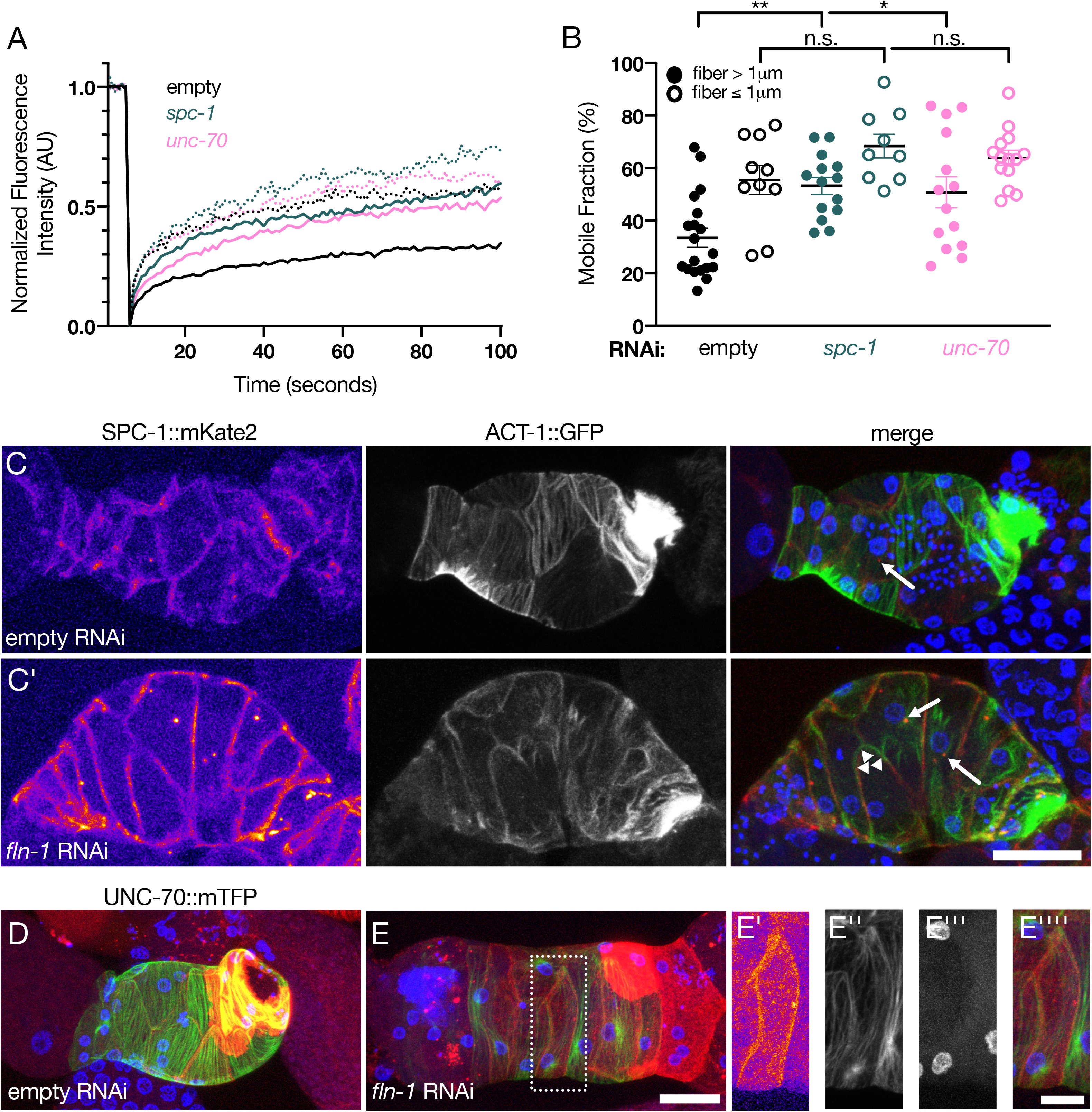
Spectrin stabilizes peripheral actin bundles in *fln-1* animals. (A) Normalized fluorescence intensity of bleach and recovery curves of actin in *fln-1* animals treated with empty, *spc-1*, or *unc-70* RNAi for large (>1 μm, solid lines) or small (≤1 μm, dotted lines) fibers. (B) Mobile fraction of actin recovery after bleach in *fln-1* animals treated with empty, *spc-1*, or *unc-70* RNAi for large (>1 μm, solid circles) or small (≤1 μm, open circles) fibers. Each point represents a measurement from single bleach and recovery curve. Error bars are SEM. Ordinary one-way ANOVA with Tukey’s multiple comparison: *** p ≤ 0.001, * p ≤ 0.05. (C-C’) SPC-1∷mKate2 localization (Fire LUT, first panel; red in merge) with respect to ACT-1∷GFP with and without *fln-1*. Arrows indicated perinuclear SPC-1 dots. Arrowheads indicated thick peripheral actin bundles associated with junctional spectrin. Scale bar is 20 μm. (E) UNC-70 tagged with mTFP (red), F-actin labeled with moeABD∷mCherry (green), and nuclei labeled with DAPI (blue) with (D) and without (E) *fln-1*. Scale bar is 20 μm. (E-E’’’’) Dotted box indicates a cell with UNC-70 (E’, fire LUT, red in merge), actin (E’’, green in merge), and nuclei (E’’’, blue in merge) where actin is localized along UNC-70 rich cell junctions (E’’’’ merge). Scale bar is 10 μm.

In animals treated with either empty or *fln-1* RNAi, SPC-1∷mKate2 is localized primarily to cell junctions as described previously (Figure 7C-C’) (Wirshing & Cram, 2018). In spermathecae in which *fln-1* is depleted, some of the large fibers appear to localize to the cell periphery with SPC-1 (Figure 7C’). These data suggest that spectrin is involved in stabilizing thick peripheral bundles in animals lacking *fln-1*. UNC-70 localization along cell junctions was similar to that of SPC-1 (Figure 7D). Treatment with *fln-1* RNAi results in colocalization of the thick peripheral actin bundles and peripheral UNC-70 (Figure 7E-E’’’’). These results suggest the SPC-1/UNC-70 spectrin complex localizes to and is required for thick peripheral bundle stability in *fln-1* animals.

### Loss of filamin results in mislocalization and disrupted morphology of several organelles

The actin cytoskeleton is intimately connected to and can govern the morphology and distribution of organelles and membranes (Breuer et al., 2017; Deneke et al., 2019; Keeling, Flores, Dodhy, Murray, & Gavara, 2017; Korobova, Ramabhadran, & Higgs, 2013; Köster et al., 2016; Ueda et al., 2010). Given that the actomyosin cytoskeleton was so severely disrupted in animals lacking *fln-1*, we hypothesized that other components of the cell would be mislocalized. We imaged nuclei, mitochondria, and the endoplasmic reticulum to assess global cellular organization. We first looked at nuclear position by imaging excised gonads of animals expressing GFP labeled membranes (MmPLCδ∷GFP), and staining for F-actin and nuclei with phalloidin and DAPI, respectively (Figure 8A-A’). Nuclei in the spermathecal cells are positioned so that they are touching or nearby (0.37 ± 0.11 μm) the plasma membrane (Figure 8B, C). In *fln-1(RNAi)*, the nuclei that are not touching the cell edge are positioned further away from the edge (0.97 ± 0.12 μm) (Figure 8B,C). We next observed the distribution of the mitochondria and the endoplasmic reticulum in the spermatheca. We used a fragment of TOMM-20∷GFP to label mitochondria and observed striking differences between animals fed either control or *fln-1* RNAi. Normally mitochondria are distributed across the cells, however without filamin, they clump near the nucleus and are absent from other areas of the cell (Figure 8 D-E). Similarly the endoplasmic reticulum, labeled endogenously with GFP∷SEC-61.B, is also mislocalized and unevenly distributed to clumps near the cell periphery rather than being evenly distributed (Figure 8F-G). We conclude that filamin either directly or indirectly through its effects on actin network organization, functions to maintain global cellular organization of organelle morphology and distribution.

**Figure 8:**
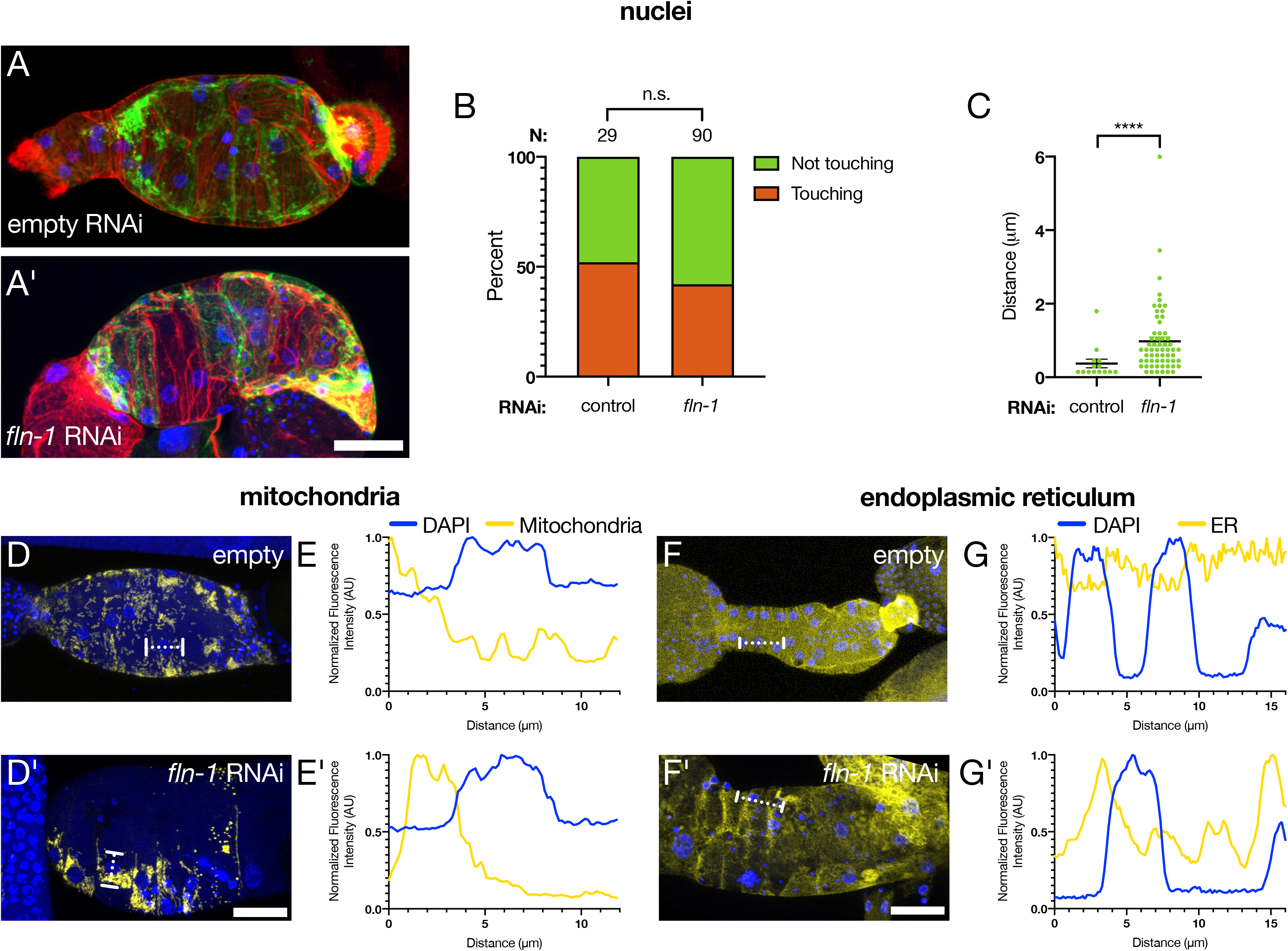
Filamin is required for correct positioning, size, and distribution of organelles. (A) Spermathecae of excised gonads expressing MmPLCδ∷GFP to label membrane (green), were stained with phalloidin to label F-actin (red), and DAPI to label nuclei (blue). Knockdown of *fln-1* using RNAi results in redistribution of nuclei with respect to cell peripheries (A’) Scale bar is 20 μm. (B) There is no difference in the percent of nuclei that are either touching, or not touching a cell edge. N represents the number of nuclei analyzed. Fisher’s exact test: ns. (C) Of the nuclei that are not touching the cell periphery, the distance between the nuclear edge, and the cell membrane is increased without *fln-1*. Each point represents a nucleus measured. Control (empty vector) RNAi = 14 cells (4 animals); *fln-1* RNAi = 61 cells (14 animals). Mann Whitney test: **** p < 0.0001. (D-D’) Mitochondria labeled with an fragment of TOMM-20 fused to GFP (yellow) and nuclei labeled with DAPI (blue) show that animals treated with *fln-1* RNAi have clumps of mitochondria near nuclei, whereas empty vector results in an evenly distributed mitochondrial network. Scale bar is 20 μm. Line scans, indicated with a dotted line in E-E’, are plotted for control and *fln-1* RNAi treated spermathecal cells in D-D’. Mitochondria are less evenly distributed, and tightly associated with nuclei in *fln-1* animals. (F-F’) The endoplasmic reticulum is labeled with an endogenous eGFP∷Sec-61B tag (yellow) and have been excised and stained with DAPI (blue) Scale bar is 20 μm. Animals fed *fln-1* RNAi have less evenly distributed endoplasmic reticulum than empty vector fed animals, quantified in G-G’ with a line scan (dotted line in F-F’).

## Discussion

In the spermatheca, myosin activity (Kelley et al., 2018; Wirshing & Cram, 2017) and actin interacting proteins (Deng et al., 2007; Kovacevic & Cram, 2010; Wirshing & Cram, 2018) organize the actomyosin network into basal stress fiber-like bundles that form during contraction and are required for productive contractility of the tissue. We show that, along with playing a crucial role in organizing the actin cytoskeleton, filamin/FLN-1 is required for correct localization and distribution of nonmuscle myosin. Loss of *fln-1* results in the formation of myosin foci, which are localized to the nucleus by the LINC complex components ANC-1 and UNC-84. FLN-1 is also required for the anchorage of actin bundles to the basal surface. We show that peripheral actin bundles do not interact with myosin, but rather colocalize with and are stabilized by actin interacting proteins such as spectrin. Without FLN-1, the cells undergo dramatic rearrangements of not only the cytoskeleton and associated proteins, but also of several organelles. Therefore, filamin/FLN-1 is a key organizer of the cytoplasm in spermathecal cells.

In the mature spermatheca, myosin colocalizes with actin in contractile bundles (Wirshing & Cram, 2017). Therefore, it was surprising that without FLN-1, myosin primarily associates with actin near the nucleus. We speculate that spermathecal myosin may have a higher affinity for actin cables that are under tension. If so, when filamin is lost and tension decreases, myosin might then relocalize to other binding sites. Myosin can be recruited to and stabilized at sites of tension in the epithelia and wing disc of *Drosophila* embryos (Duda et al., 2019; Fernandez-Gonzalez, de Matos Simones, Roper, Eaton, & Zallen, 2009; Kobb, Zulueta-Coarasa, & Fernandez-Gonzalez, 2017). Myosin has also been shown to be tension sensitive during cytokinesis in Dictyostelium (Luo et al., 2012; Ren et al., 2009). Several groups have also suggested that myosin prefers actin filaments that are in a stretched conformation (Shutova et al., 2012; Uyeda, Iwadate, Umeki, Nagasaki, & Yumura, 2011). Therefore in this system, it may be that there is a redistribution of forces without *fln-1*, either directly due to a loss of actin fiber anchorage or indirectly through an unidentified binding partner.

Without filamin, contraction of the basal actomyosin network results in the rapid accumulation of myosin foci near the nucleus. The LINC complex is required to mediate the proximity of the *fln-1* actomyosin foci to the nucleus. The LINC complex is comprised of a SUN-domain containing protein that spans the inner nuclear membrane and a KASH-domain containing protein that spans the outer nuclear membrane and can bind components of the cytoskeleton (Starr & Fridolfsson, 2010). We found that UNC-84, the only SUN-domain protein in *C. elegans* (Malone, Fixsen, Horvitz, & Han, 1999), and ANC-1, an actin binding KASH-domain protein (Starr & Han, 2002), are required for the nuclear localization of the actomyosin foci in *fln-1* animals. This suggests that perinuclear actin may be locally organized and possibly anchored to the global cytoskeleton. We hypothesize that loss of *fln-1*, which normally functions to keep actin anchored, results in ‘free’ actomyosin fibers that can be snared by the LINC complex at the nucleus. This perinuclear capture results in increasingly distinct puncta as the myosin contracts.

Our FRAP data reveals differences in actin dynamics depending on the size of bundles in *fln-1* mutants. Smaller fibers have an increased mobile fraction of recovery compared to larger fibers. Altering the stability of actin bundles, either through chemical perturbation, and under- or over-expression of actin binding proteins can change the recovery of actin in FRAP experiments (Bae, Sung, Cho, & Song, 2012; Lorente, Syriani, & Morales, 2014; Pappas, Krieg, & Gregorio, 2010; Zweig & Singer, 1979). In the spermatheca, the mobile fraction of small fibers was around 60%, which is consistent with the mobile fractions of actin recovery seen in the leading edge of lamellipodia in highly motile cancer cells during migration (Lai et al., 2008; Lorente et al., 2014). This suggests that without *fln-1*, small fibers are highly dynamic, and likely undergoing polymerization, depolymerization, and reorganization. In contrast, large fibers have a significantly smaller mobile fraction. This reduction in dynamics could be the result of increased bundling or changes in polymerization or depolymerization rates. We found that the increased stability of the large actin fibers depends on the expression of several actin-binding proteins. These include PLST-1/plastin and SPC-1/spectrin, proteins that bind and organize actin (Ding et al., 2017; Norman & Moerman, 2002; Otto, 1994; Shinomiya, 2012; Stevenson, Veltman, & Machesky, 2012; Wirshing & Cram, 2018), GSNL-1/gelsolin and VILN-1/villin, which can influence stability through bundling, severing, and capping actin (Klaavuniemi, Yamashiro, & Ono, 2008; Meng et al., 2015; S. Ono, 2014; Otto, 1994; Stevenson et al., 2012), and FHOD-1/formin, an actin polymerizing protein (Mi-Mi, Votra, Kemphues, Bretscher, & Pruyne, 2012; Stevenson et al., 2012; Vanneste, Pruyne, & Mains, 2013). Filamin binding to actin fibers may sterically restrict protein-actin interactions such that loss of filamin results in increased availability of binding sites or substrates for these normally excluded proteins. This screen provides a new method for identifying redundant actin bundling or stabilizing proteins, and could be expanded to include additional factors. For example, a recent publication has revealed a noncanonical role for the GTPase dynamin in bundling actin filaments (Zhang et al., 2020). Using the *fln-1* background for screens may prove useful in identifying and understanding the role for novel bundlers, like dynamin, in organizing the cytoskeletal network in vivo.

Several proteins, including spectrin and filamin, have well defined actin organizing patterns in terms of distance and angles between actin fibers (B. Han, Zhou, Xia, & Zhuang, 2017; Holley & Ashmore, 1990; Nakamura, Osborn, Hartemink, Hartwig, & Stossel, 2007; Popowicz et al., 2004; Unsain, Stefani, & Cáceres, 2018). Filamin is often studied in the context of the branched actin networks within lamellipodia where it crosslinks orthogonal actin fibers (Flanagan et al., 2001; Kumar et al., 2019; Nakamura et al., 2007). In the *C. elegans* spermatheca, the actin appears to be organized in parallel stress fiber-like actomyosin bundles rather than a meshy actin network. Recent results suggest that filamin can also localize to and act to stabilize actin bundles (Kumar et al., 2019). However, it is also possible that the spermathecal structures instead consist of disordered branched networks or disordered bundles of actin and myosin (Ennomani et al., 2016; Kassianidou, Brand, Schwarz, & Kumar, 2017; Kumar et al., 2019), which can not be distinguished from parallel bundles using traditional light microscopy. Disordered bundles or fibers with more complex geometries can produce contractions and transmit force over large distances (Ennomani et al., 2016; Kassianidou et al., 2017; Kumar et al., 2019), something required of the spermathecal basal actomyosin fibers. In branched networks or disordered bundles, filamin could be acting as it usually does to crosslink actin fibers orthogonally (Nakamura et al., 2007). If this is the case, the large organized fibers we see in wild type animals may actually be the sum of short, disordered actin fibers organized by filamin and other actin-binding proteins as identified in our FRAP screen.

We have previously published that spectrin is required to produce and maintain basal actin stress fibers in the spermatheca and is required for proper contractility of the tissue (Wirshing & Cram, 2018). Here we describe a role for spectrin in stabilizing the large, apicalized peripheral bundles that form in *fln-1* mutants. This data supports our previously published data that shows that in addition to being basally localized, SPC-1/α-spectrin and UNC-70/β-spectrin localize to the apico-lateral junctions and stabilize peripheral bundles (Wirshing & Cram, 2018). We also find that SPC-1 localizes to the perinuclear area in wild type and this localization is increased in *fln-1* animals. Nonerythroid αII-spectrin is a well-characterized spectrin subunit that has several nuclear partners including emerin (Holaska & Wilson, 2007), which in turn complexes with lamin, SUN2, and other nuclear proteins (Holaska & Wilson, 2007; Lambert, 2018; Zhong, Wilson, & Dahl, 2010). However, differences include SPC-1/α-spectrin localization to the outside of the outer nuclear membrane, indicating that this may be a newly discovered role for spectrin in the spermatheca. Understanding how filamin, spectrin, and the LINC complex coordinate to organize the cytoskeleton and nuclear-associated actin will be an exciting new direction of study, where we may better understand the role of the nucleus as a mechanical sensor and organizer of the cytoskeleton.

Loss of filamin/FLN-1 not only altered cytoskeletal organization, but also resulted in dramatic rearrangement of organelles. Filamin could be directly interacting with an organelle, or its associated cytoskeleton, to shape and position it, as seen in nuclei during Drosophila nurse cell dumping (Huelsmann, Ylänne, & Brown, 2013). Or, filamin’s effect on organelle morphology and placement could be indirect, through one of its many known protein interactors (Feng & Walsh, 2004). For instance, several of the kinases known to interact with filamin such as PKA (Jay, Garcia, & de la Luz Ibarra, 2004) and PKC (Tigges, Koch, Wissing, Jockusch, & Ziegler, 2003), influence the mitochondrial network and activity (Lucero, Suarez, & Chambers, 2019). Lastly, the reorganization of organelle position and morphology may be through the indirect effects of cytoskeletal disruption in *fln-1* mutants. The cytoskeleton interacts with and organizes several organelles in terms of localization and morphology including nuclei (Deneke et al., 2019; Keeling et al., 2017; Starr & Han, 2002), mitochondria (Korobova et al., 2013), and the endoplasmic reticulum (Ueda et al., 2010), along with other organelles and membranes (Breuer et al., 2017; Köster et al., 2016).

Because humans have three filamins (Stossel et al., 2001), while *C. elegans* has two, FLN-1 and FLN-2 (DeMaso et al., 2011), *C. elegans* provides a simplified system to explore the role of filamin in organizing the actomyosin cytoskeleton. FLN-1 is the only filamin expressed in the spermatheca. This may explain why the phenotypes we observed are more dramatic than the loss of filamin phenotypes seen in other systems. Various familial filaminopathies are associated with filamin A, B, and C in humans, although the molecular mechanisms behind these disorders are not well understood (Baudier, Jenkins, & Robertson, 2018; Duff et al., 2011; Ferrer & Olivé, 2008; Kley et al., 2007; Razinia et al., 2012; Vorgerd et al., 2005; Wade, Halliday, Jenkins, O’Neill, & Robertson, 2020). Our work also suggests that in addition to the known roles of filamin as an actin crosslinker and signaling scaffold, loss of filamin in non-redundant systems can result in global rearrangement of the cell cytoskeleton and organelles.

## Abbreviations

FRAP: fluorescence recovery after photobleaching
GFP: green fluorescent protein
KASH: Klarsicht/ANC-1/Syne homology
LINC: linker of nucleoskeleton and cytoskeleton
moeABD: moesin actin binding domain
RNAi: RNA interference
sp-ut: spermatheca-uterine
SUN: Sad1/UNC-84
WT: wild type

## Acknowledgements

We would like to thank Daniel Starr and Hongyan Hao for providing the endogenously labeled GFP∷ANC-1 strain, and for helpful discussions. We would also like to thank David Calderwood and Ronen Zaidel-Bar for helpful discussions. Several strains used in this study were provided by the Caenorhabditis Genetics Center, which is funded through the National Center for Research Resources, National Institutes of Health. This work is funding by the National Science Foundation/Molecular and Cellular Biosciences–U.S. Israel Binational Science Foundation award (1816640) to E.J.C. and R.Z.-B.

## Materials and Methods

### Nematode maintenance and strain generation

All animals were grown at 23°C on Nematode Growth Media (NGM) (0.107 M NaCl, 0.25% wt/vol Peptone, 1.7% wt/vol BD Bacto-Agar, 2.5 mM KPO_4_, 0.5% Nyastatin, 0.1 mM CaCl_2_, 0.1 mM MgSO_4_, 0.5% wt/vol cholesterol) and fed OP50 *Escherichia coli* (Hope, 1999). For experiments, animals were synchronized using an “egg prep”, where gravid hermaphrodites were lysed in an alkaline hypochlorite solution, and washed several times in M9 buffer solution (22 mM KH_2_PO_4_, 42 mM NaHPO_4_, 86 mM NaCl, and 1 mM MgSO_4_) (Hope, 1999). Embryos were then pipetted onto new plates and allowed to grow at 23°C for 52-72 hours, depending on the strain and experiment. All extrachromosomal arrays were injected as described previously (Mello, Kramer, Stinchcomb, & Ambros, 1991). UN1911 and UN1913 were made by crossing UN0810 *fln-1(tm545)* (Kovacevic & Cram, 2010) with UN1654 [*GFP∷NMY-1;moeABD∷mCherry*] and UN1502 [ACT-1∷GFP], respectively. (Wirshing & Cram, 2017). UN1953 was generating by crossing UN0810 *fln-1(tm545)* (Kovacevic & Cram, 2010) with EU573 *orEx2 [mlc-4p∷mlc-4∷GFP∷unc-54 3’UTR + rol-6(su1006)]* (K. Ono & Ono, 2016). WBM875 [*Y38F2AR.9(wbm3[eGFP∷Sec61b])IV]* was a gift from the Mair lab and used to label the endoplasmic reticulum. See Supplementary Table 2 for a complete list of strains used.

### RNAi Treatment

The RNAi feeding method was used as described previously (Kovacevic & Cram, 2010). HT115(DE3) bacteria with a double stranded construct in the L4440 backbone were grown shaking overnight in Luria broth at 37°C, then the bacteria were seeded onto NGM plates with 25 μg/ml carbenicillin and 1 mM isopropylthio-β-galactoside (IPTG).

### Fluorescent Imaging

Gonads were excised from age-synchronized animals using 23-guage needles in phosphate-buffered saline (PBS) then transferred to a 15 ml glass conical tube. Tissues were fixed in 1.85% formaldehyde in PBS for 25 minutes at room temperature, washed twice in PBS, then permeablized in PBS + 0.1% Trition X-100 (PBST) for 15 minutes. Gonads were incubated overnight at 4°C with 1 U/ml Texas Red-X Phalloidin (ThermoFisher Scientific) in staining solution (1% BSA, 0.05% Triton X-100 in PBS. To visualize the nucleus, 4’,6-diamidino-2-phenylindole (DAPI; Sigma-Aldrich) was added at a final concentration of 20 ng/μl. Gonads were then washed twice with PBS at room temperature, and mounted on a 2% agarose pad. Coverslips were sealed with nail polish prior to imaging. To visualize nuclei in the animals with labeled actin and myosin (moeABD∷mCherry; GFP∷NMY-1), gonads were dissected fixed and rinsed as described above, then stained with DAPI at a concentration of 100 ng/μl in staining solution all in a watchglass. Gonads were stained for 30 minutes, covered, at room temperature, then washed twice in PBS, and mounted as described above. Imaging of organelles was done with three-channel imaging of excised and fixed gonads stained Texas Red-X Phalloidin and DAPI as described above, excited with a 405-nm (DAPI), 488-nm (GFP), and 594-nm laser, switching each line with a pinhole of 1 AU for each laser line. Z-intervals were 0.32 μm.

For ovulation movies animals were synchronized using the egg prep protocol described above. Animals were immobilized on a 5% agarose pad with a 1:1:1.5 ratio of 0.01% tetramisole and 0.1% tricaine solution in M9 buffer, M9 buffer, and 0.05 μm Polybead microspheres (Polysciences) and only first ovulations were imaged. Imaging was done on a Zeiss LSM 710 confocal using Zen software and a Plan-Apochromat 63x/1.40 oil objective lens. GFP and mCherry were excited with 488 and 561 nm lasers, respectively. DAPI and Texas Red-X dyes were excited at 405 and 594 nm lasers, respectively. Time-lapse imaging was performed by capturing 30 (two channels) or 40 (single channel) z-slices at 12 and 15-second intervals, respectively. Still images were captured at 0.32 μm (images including DAPI) or 0.46 μm (phalloidin only) intervals with each slice being averaged two to four times. FRAP was performed using the 488 nm laser and captured at 1 second intervals and a 1.5 AU pinhole (1.1 μm section). After 5 slices, a 31×12 pixel region (~3.1×1.2 μm) region was bleached using the 405-nm and 488-nm lasers at 2% and 100%, respectively using a slow scan speed (14.35 μsec/pixel), then imaged for recover for an additional 95 slices.

### Image Analysis

All image analysis was performed using ImageJ software. Movies of actin and myosin (moeABD∷mCherry; GFP∷NMY-1) had the background subtracted using the rolling ball method with a radius of 50 pixels. Both GFP∷ACT-1 and moeABD∷mCherry; GFP∷NMY-1 movies had their intensities adjusted for easier visualization. Line scans of actin and myosin intensity with respect to distance to the nucleus, was performed by drawing a 10 pixel-wide line across the nucleus and myosin foci using a maximum intensity projection images. Line intensities were plotted, and the average distance per spermatheca between the peak DAPI signal (edge of nucleus) and peak myosin intensity was the distance between the foci and nucleus. For image analysis of the moeABD∷mCherry;DEB-1∷3xGFP animals, a line scan using a 10 pixel-wide line was made across the width of the cell, normalized to the background intensity of each channel, and plotted to show nuclear association of vinculin colocalized with the nuclear pool of actin. To quantify the intensity of DEB-1 within and along the perimeter of cells, the average intensity within an ROI drawn along the cell boundaries and along (5pixel wide line average) was calculated. The values graphed are the average intensities along the perimeter of each cell divided by the area within each cell. FRAP data was analyzed as described previously (Kelley et al., 2018), using a Jython script for ImageJ. In order to quantify nuclear positioning in control and *fln-1* RNAi treated animals, single slices of each z-stack for a given nuclei were analyzed. A 5-pixel wide line was drawn across the nearest membrane/cell edge of the nucleus and the distance between the edge of the nucleus and the edge of the cell membrane was quantified. If the number was >0 that nucleus was categorized as not touching the cell membrane. For Figure 8C, the distance between nucleus and cell membrane was plotted only for those nuclei that were not touching the cell edge. To quantify distance between nuclei and either mitochondria or endoplasmic reticulum, a 50 or 25 pixel line, respectively, was used to generate a line intensity profile in ImageJ and plotted using Graphpad Prism,

### Statistical Analysis

The Pearson’s correlation coefficient was calculated using Matlab. Briefly, for each frame of the movie, the image was cropped so that the ROI contained the central cells of the spermathecal bag (removing the sp-ut valve, which is only labeled with actin∷GFP). Each image was smoothed using the ‘smoothdata’ function, with a moving average (‘movmean’) over a window of 3 pixels, then the correlation between the red and green channels was calculated for each frame and plotted over time. All other statistics were performed using GraphPad Prism 8 Software. Unpaired t test was used to compare two groups. For more than two groups, an ordinary one-way ANOVA with a Tukey’s multiple comparisons test was used to compare the mean of each group to every other group in the data set. For all data, symbols are represented as follows: p > 0.05, * p ≤ 0.05, ** p ≤ 0.01, *** p ≤ 0.001, **** p ≤ 0.0001.

**Supplemental Table 1.**

| Gene | Protein | Function | Role in actin organization | Expression | Sources |
| --- | --- | --- | --- | --- | --- |
| ***alp-1*** | ALP/<br>Enigma | actin organization in body wall muscle, neuron regeneration | Predicted actin binding activity, localizes to dense bodies/Z-discs, involved in Rho kinase/LET-502 activation | Body wall muscle, pharynx, hypodermis, hypodermal seam cells, pharynx | (H.-F. Han & Beckerle, 2009; McKeown, Han, & Beckerle, 2006; Shimizu, Pastuhov, Hanafusa, Matsumoto, & Hisamoto, 2018) |
| ***atn-1*** | alpha actinin | Sarcomere organization and body movement | Has two actin binding (CH) domains, localizes to dense bodies/Z-discs, thin filaments, and cytoplasmic actin filaments | Body wall muscle, some neurons, | (Barstead, Kleiman, & Waterston, 1991; Hresko, Williams, & Waterston, 1994; Moulder et al., 2010; S. Ono, 2014; Otto, 1994; Stevenson et al., 2012; Winkelman et al., 2016) |
| ***fhod-1*** | formin | Embryonic morphogenesis | Localizes to the dense bodies/Z-lines | Spermatheca, body wall muscle, pharynx, vulva, embryo | (Mi-Mi et al., 2012; Stevenson et al., 2012; Vanneste et al., 2013) |
| ***fli-1*** | flightless-I | Helps establish A/P polarity in the single cell embryo, required for neuron cell and axon migration, germline development, ovulation, | Predicted actin filament binding activity. Organizes actin in embryos, spermathecae, body wall muscle, and germline | Spermatheca, sp-ut valve, distal tip cell, body wall muscle, pharynx, vulva, proctodeum muscle (males), neurons, embryos | (Campbell et al., 1993; Deng et al., 2007; Goshima et al., 1999) |
| ***gsnl-1*** | gelsolin | Synapse plasticity and remodeling, | Actin severing and capping protein, binds actin filaments | Body wall muscle, neurons | (Fox et al., 2007; Klaavuniemi et al., 2008; Meng et al., 2015; S. Ono, 2014; Otto, 1994; Stevenson et al., 2012) |
| ***inft-2*** | formin | Tubulogenesis of the excretory cell, mobility of the posterior body | Promotes apical F-actin accumulation | Excretory cell, spermatheca | (Mi-Mi et al., 2012; Shaye & Greenwald, 2016; Stevenson et al., 2012; Wirshing & Cram, 2017) |
| ***plst-1*** | plastin | Embryonic polarization and cytokineses | Actin bundling and crosslinking | Socket cells in amphids and phasmids, pharyngeal-intestinal valve, intestine, distal gonad, vulva, and intestinal-rectal valve, embryo | (Ding et al., 2017; Otto, 1994; Shinomiya, 2012; Stevenson et al., 2012) |
| ***spc-1*** | alpha spectrin | Embryonic morphogenesis, spermathecal contractility | Interacts with beta spectrin to stabilize cortical actin meshwork, helps designate actin bundle spacing, distribution,<br>and orientation | Spermatheca, sheath, spermatheca-uterine valve, embryos, neurons | (Norman & Moerman, 2002; Stevenson et al., 2012; Wirshing & Cram, 2018) |
| ***unc-115*** |  | Axon guidance,<br>neuronal<br>morphogenesis | Actin binding, contains<br>villin headpiece<br>domain | Neurons | (Lundquist, Herman,<br>Shaw, & Bargmann,<br>1998; Yang &<br>Lundquist, 2005) |
| ***viln-1*** | supervillin |  | Predicted actin<br>filament binding |  | (Otto, 1994;<br>Stevenson et al.,<br>2012) |
| ***zyx-1*** | zyxin | LIM-domain<br>containing protein<br>that regulates<br>recruitment of<br>proteins to dense<br>bodies, body<br>bending, interacts<br>with germline RNA<br>helicase | Localizes to dense<br>bodies, M-line, and<br>nucleus of body wall<br>muscle | Spermatheca,<br>Body wall<br>muscle, some<br>neurons | (Lecroisey et al.,<br>2013, 2008;<br>Nahabedian, Qadota,<br>Stirman, Lu, &<br>Benian, 2012; Smith<br>et al., 2002) |

**Supplemental Table 2.**
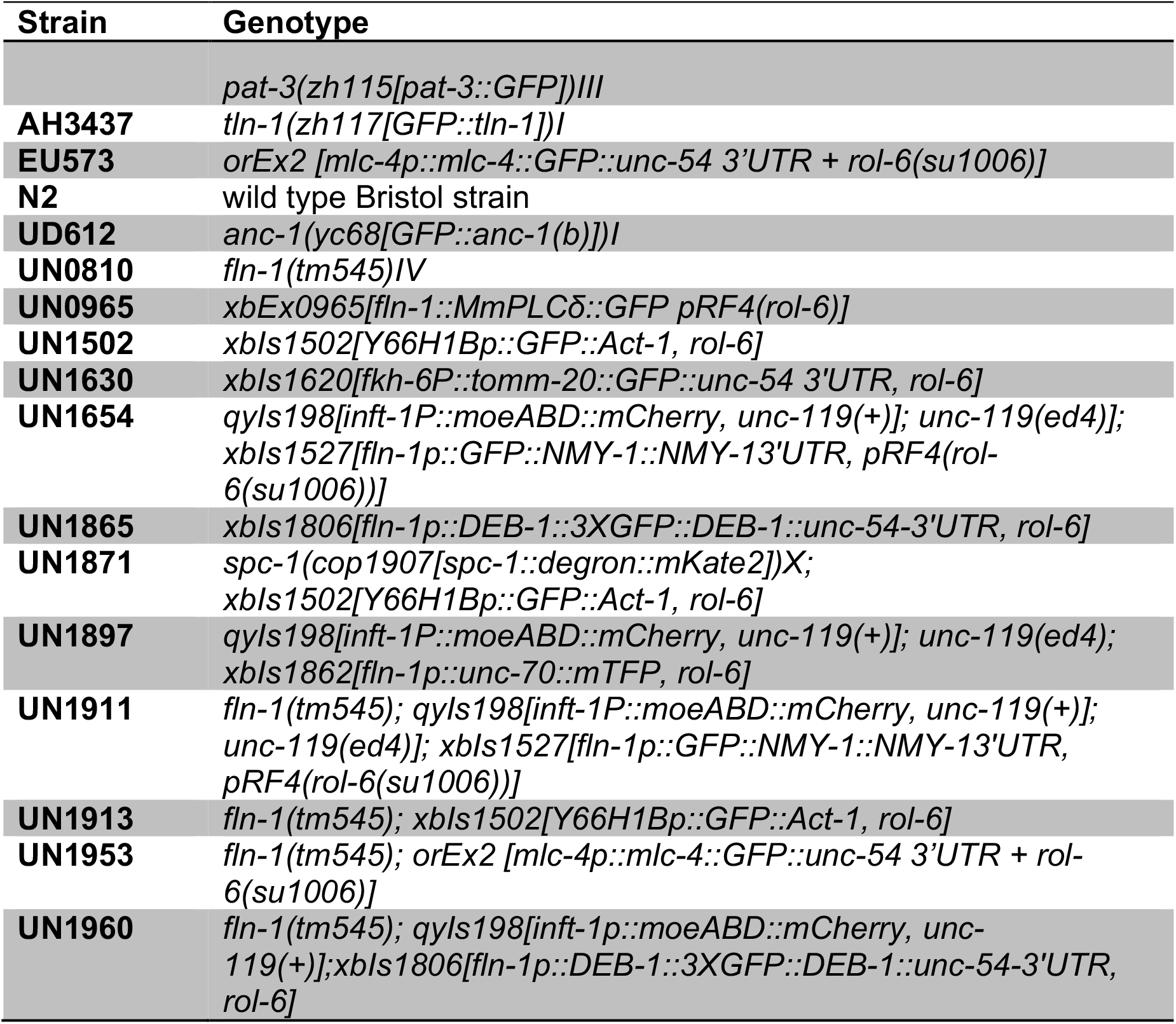
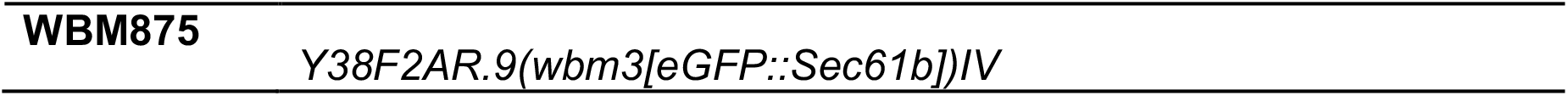

---

**Supplemental Figure 1: Filamin is required for colocalization of actin and myosin during contraction**

Pearson correlation coefficient plots for each frame of wild type (A) and *fln-1(tm545)* (B) time lapse movies imaged. Calculations begin when the egg begins to enter the spermatheca. Dotted lines labeled “a” indicate when the egg is fully in the spermatheca, and “b” to indicate the start of exit for wild type animals.

**Supplemental Figure 2: Focal adhesion components localize to basal actin structures and to the cell periphery**

TLN-1 (green) with F-actin (red) and nuclei stained with Phalloidin and DAPI, respectively with (A-A’’ maximum intensity projection, B-B’’ cross section of box indicated in A’’) or without (C-C’’) *fln-1*. PAT-3 (green) with F-actin (red) and nuclei stained with Phalloidin and DAPI, respectively with (D-D’’ maximum intensity projection, E-E’’ cross section) or without (F-F’’) *fln-1*. Scale bar for A, C, E, and F is 20 μm, scale bar for B and E is 10 μm.

**Supplementary Figure 3: Several genes regulate the stability of large *fln-1* actin bundles**

(A-L) Normalized fluorescence intensity of the bleach and recovery curves of *fln-1* animals treated with RNAi for large (>1 μm, solid lines) or small (≤1 μm, dotted lines) fibers. Animals treated with control RNAi are black in all traces. (A’-L’) Quantification of the mobile fraction for *fln-1* animals treated with empty (black dots in all graphs) or experimental RNAi for large (>1 μm, solid circles) or small (≤1 μm, open circles) fibers. Each point represents a single bleached fiber recovery. For each condition, only matched sizes were compared using an unpaired t test: * p ≤ 0.05, ** p ≤ 0.01. Error bars are SEM.

## Supplemental Movies

**Supplemental Movie 1: Actin becomes aligned during the first ovulation.** Maximum intensity projection of a 4D confocal movie of a wild type first ovulation expressing ACT-1∷GFP. The when the oocyte enters (~2:48), the actin is initially unorganized, but as the tissue begins to contract and the sp-ut valve opens (~8:24) the fibers become aligned. The newly fertilized embryo is then pushed into the uterus and the spermatheca returns to the unoccupied conformation. Z-stacks were imaged at 14-second intervals and played back at 30 frames per second. Scale bar is 20 μm.

**Supplemental Movie 2: FLN-1 is required for proper development of the actin network.** Maximum intensity projection of a 4D confocal movie of a *fln-1(tm545)* first ovulation expressing ACT-1∷GFP. The oocyte enters the spermatheca (~2:30), and initially the actin is unorganized. The spermathecal tissue does not contract to push the egg into the uterus, rather actin appears to be “pulled” into a point in each cell (~27:30) then the ACT-∷GFP signal disperses throughout the cell. Z-stacks were imaged at 15-second intervals and played back at 30 frames per second. Scale bar is 20 μm.

**Supplemental Movie 3: Actin and myosin colocalize during contraction.** Maximum intensity projection of a 4D confocal movie of wild type first ovulation with labeled actin and myosin with moeABD∷mCherry and GFP∷NMY-1, respectively. The oocyte enters the spermatheca (~5:12), and actin and myosin colocalize as the tissue contracts to push the embryo into the uterus. Z-stacks were imaged at 12-second intervals and played back at 30 frames per second. Scale bar is 20 μm.

**Supplemental Movie 4: Loss of *fln-1* results in separation of actin and myosin.** Maximum intensity projection of a 4D confocal movie of *fln-1(tm545)* first ovulation with labeled actin and myosin with moeABD∷mCherry and GFP∷NMY-1, respectively. The oocyte enters (~5:48) then instead of contracting, myosin rich foci begin to form in each cell (begins ~23:24) until suddenly myosin and actin appear to pull away from each other drastically (~32:00). Z-stacks were imaged at 12-second intervals and played back at 30 frames per second. Scale bar is 20 μm.

