## Supplementary figures and images for "FLN-1/Filamin is required to anchor the actomyosin cytoskeleton and for global organization of sub-cellular organelles in a contractile tissue"

### Supplemental Figures

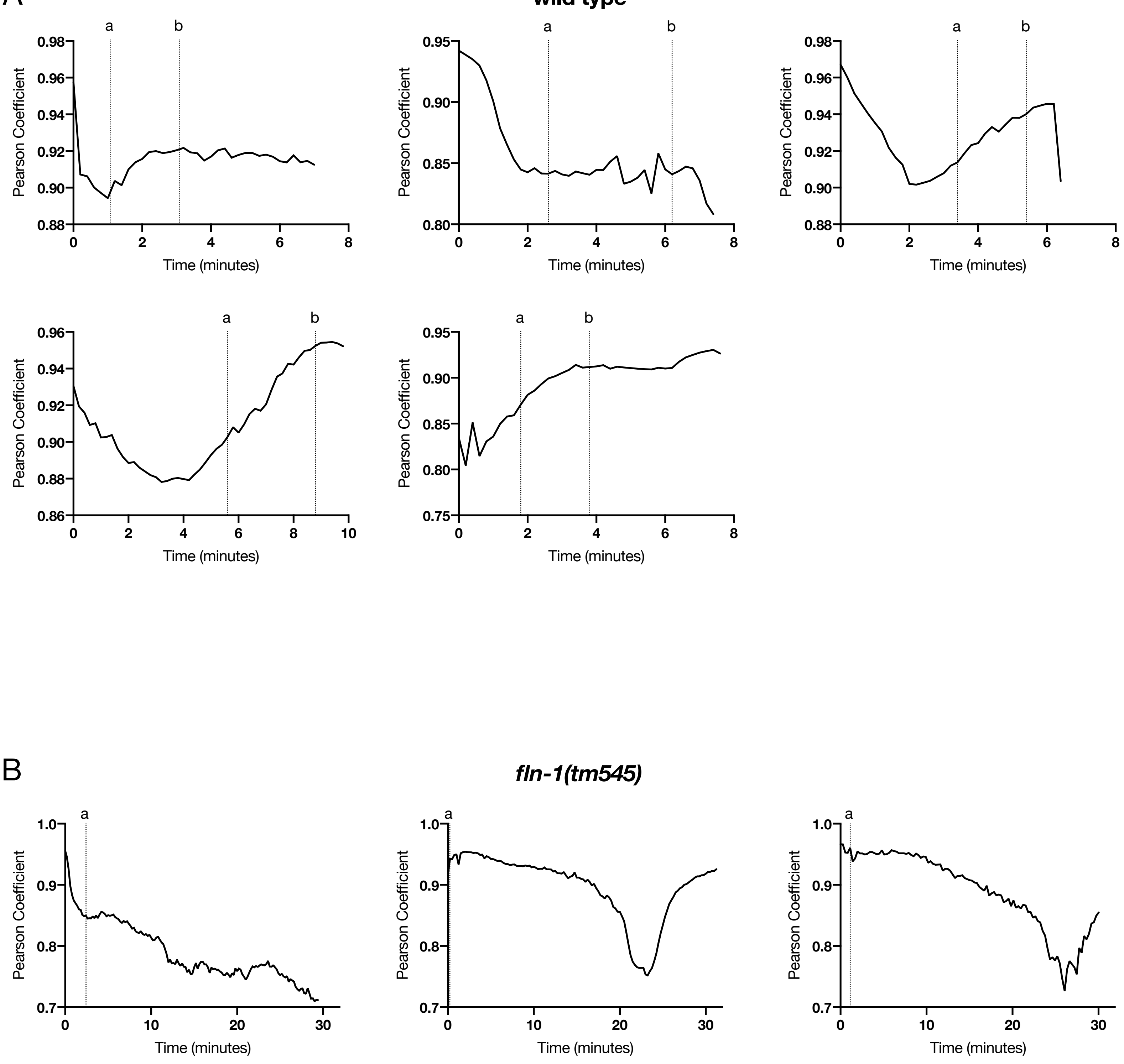

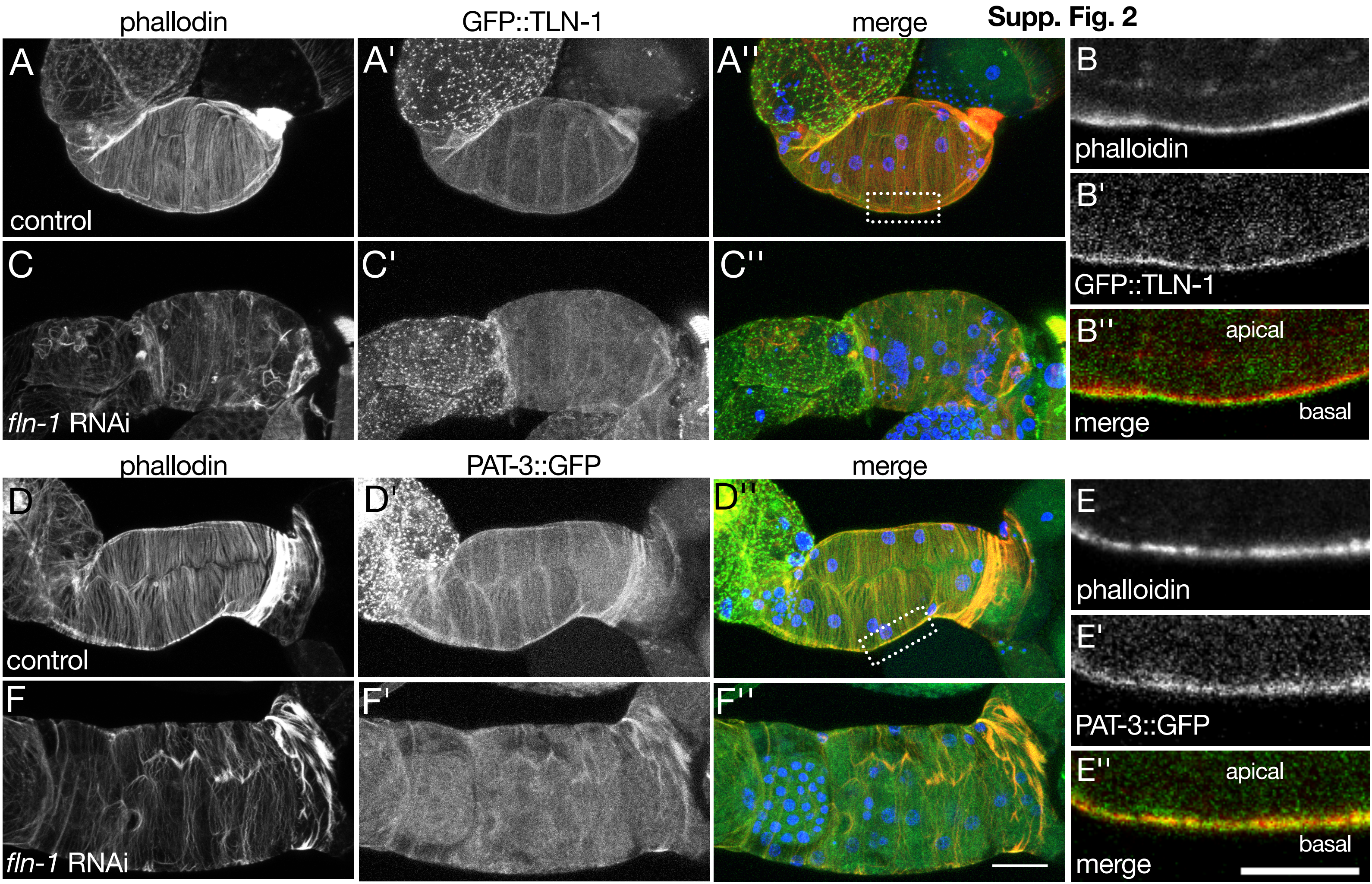

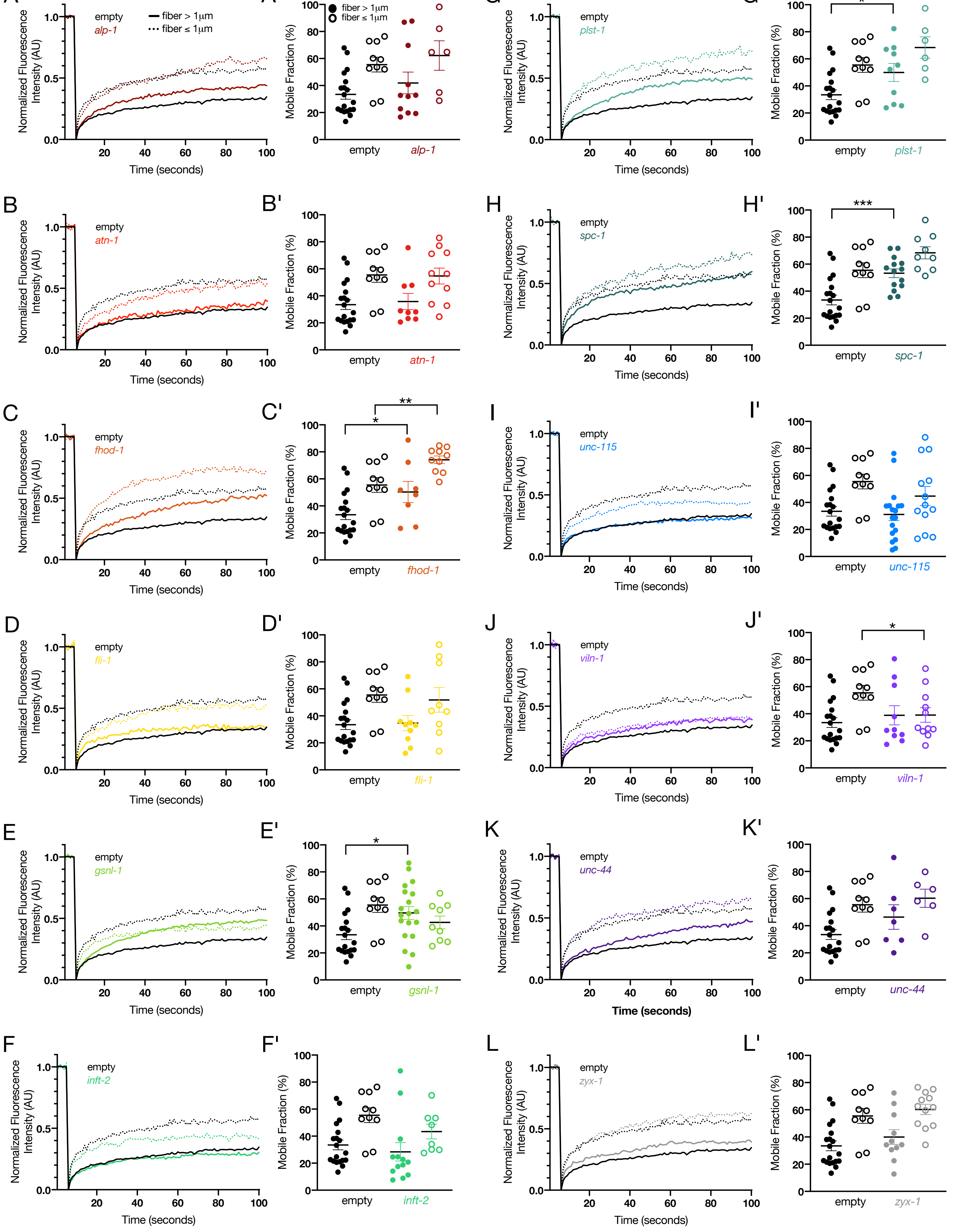
